# Conserved 5-methyluridine tRNA modification modulates ribosome translocation

**DOI:** 10.1101/2023.11.12.566704

**Authors:** Joshua D. Jones, Monika K. Franco, Mehmet Tardu, Tyler J. Smith, Laura R. Snyder, Daniel E. Eyler, Yury Polikanov, Robert T. Kennedy, Rachel O. Niederer, Kristin S. Koutmou

## Abstract

While the centrality of post-transcriptional modifications to RNA biology has long been acknowledged, the function of the vast majority of modified sites remains to be discovered. Illustrative of this, there is not yet a discrete biological role assigned for one the most highly conserved modifications, 5-methyluridine at position 54 in tRNAs (m^5^U54). Here, we uncover contributions of m^5^U54 to both tRNA maturation and protein synthesis. Our mass spectrometry analyses demonstrate that cells lacking the enzyme that installs m^5^U in the T-loop (TrmA in *E. coli*, Trm2 in *S. cerevisiae*) exhibit altered tRNA modifications patterns. Furthermore, m^5^U54 deficient tRNAs are desensitized to small molecules that prevent translocation *in vitro.* This finding is consistent with our observations that, relative to wild-type cells, *trm2*Δ cell growth and transcriptome-wide gene expression are less perturbed by translocation inhibitors. Together our data suggest a model in which m^5^U54 acts as an important modulator of tRNA maturation and translocation of the ribosome during protein synthesis.

## Introduction

Post-transcriptional modifications impact RNA structure, function, stability, and dynamics. Cells utilize these chemical modifications to control protein expression, and to ensure the speed and accuracy of the ribosome. Their significance is underscored by observations that the dysregulation of RNA modifying enzymes is linked to a myriad of pathologies including diabetes, neurological disorders, and many cancers^1–6^. While over 150 different chemical modifications exist within thousands of RNAs across all three kingdoms of life^7^, the molecular level consequences of the vast majority of modified sites remain to be discovered. Even within the most well-studied class of RNA modifications, *E. coli* and *S. cerevisiae* tRNAs, < 50% of commonly modified positions have assigned functional roles within the protein synthesis pathway.

Emblematic of this, despite six decades of study, no function is ascribed to the universally conserved 5-methyluridine modification found at position U54 in nearly all tRNA T-loops (m^5^U54) ^7–13^ (**Figure 1**). The conservation of m^5^U54 is a long standing conundrum as bacterial and eukaryotic cells lacking uracil-5-methyltransferase (Δ*trmA* in bacteria, *trm2*Δ in eukaryotes) do not exhibit a growth defect under normal laboratory conditions^14,15^. However, the ability of wild-type cells to outcompete Δ*trmA* and *trm2*Δ strains suggests that the enzymes are somehow advantageous for cellular fitness^15,16^. The origin of the fitness advantage conferred by uracil-5-methyltransferases remain enigmatic. *In vitro* translation studies using m^5^U-depleted tRNAs demonstrate that loss of m^5^U54 from tRNAs does not slow peptide elongation by the ribosome ^16–18^. m^5^U has also recently been discovered in eukaryotic mRNAs^19–22^, providing the modification a possible non-tRNA mediated avenue to influence protein synthesis. Nevertheless, the low level of m^5^U in mRNAs (at least 10-fold less than N6-methyladenosine (m^6^A) and pseudouridine (Ψ) modifications), and its minor (0 to 2-fold) effect on the rate constant for amino acid addition makes m^5^U in mRNAs unlikely to broadly enhance cellular fitness^19^.

**Figure 1.**
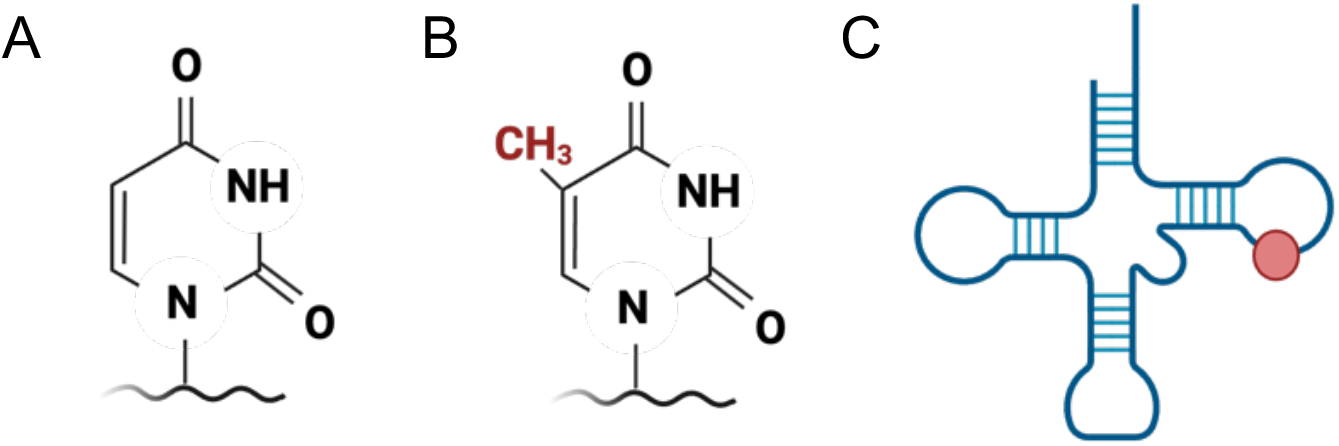
*5-methyluridine structure and location*. Structures of (**A**) uridine and (**B**) 5-methyluridine (m^5^U). (**C**) m^5^U54 is found in the T loop of tRNAs.

The most compelling evidence of a wide-spread role for tRNA uracil-5-methyltransferases is the modest destabilization of tRNAs purified from Δ*trmA* and *trm2*Δ cell lines^23^. However, it is unclear if the enzymes’ effect on tRNA stability is directly attributable to their catalysis of m^5^U54 modification, or the tRNA-folding chaperone activities reported for these enzymes ^14–16^. Further complicating matters, the “tRNA foldase” activity of TRMA does not fully account for the Δ*trmA* fitness loss relative to wild-type cells; mutational studies reveal that the ability to catalyze m^5^U54 insertion is also required to rescue cellular growth in competition assays, suggesting the modification itself enhances fitness^16^. Despite its conservation and apparent contributions to tRNA structure, the overall biological significance of the tRNA m^5^U54 modification and its contributions (if any) to protein translation are not defined.

In this work, we identify contributions of m^5^U54 to tRNA maturation, translation elongation and gene expression. Our studies reveal that *trm2*Δ cells are both desensitized to translocation inhibitors and exhibit altered tRNA modification profiles. *In vitro* translation studies support these cellular observations, demonstrating that tRNA^Phe^ purified from Δ*trmA* cells (tRNA^Phe,-m5U^) has an altered modification profile and permits the translocation of the ribosome in the presence of hygromycin B. Together, our data reveal biological roles for m^5^U54 - to promote the modification of tRNAs and modulate the speed of ribosome translocation. We find that even the subtle impact of m^5^U54 on translation has broad consequences for gene expression under cellular stress.

## RESULTS

### TRM2 impacts cell growth, transcriptome composition and reporter protein production under translational stress

The significance of RNA modifications often becomes most apparent under cellular stress when protein synthesis (and not just transcription) is particularly important for controlling the composition of the proteome^24^. With this in mind, we surveyed the impact TRM2 on cell growth under sixteen different conditions using spot plating assays. We compared the growth of wildtype and *trm2*Δ *S. cerevisiae* (BY4742) cells at varied temperature (22°C, 30°C, 37°C), carbon source (glucose, sucrose, galactose), pH (4.5, 6.8, 8.5), salt concentration (NaCl, MgSO_4_), and proteasome (MG132) and translation inhibitors (hygromycin B, cycloheximide, puromycin, paromomycin) **(Extended Data Figure 1)**. Wildtype and *trm2*Δ *S. cerevisiae* grew similarly regardless of temperature, carbon source, pH, MgSO_4_ concentration, or the presence of a proteasome inhibitor, while *trm2*Δ exhibited subtly enhanced growth over wildtype under 1 M NaCl salt stress.

By comparison, three translational inhibitors, hygromycin B, cycloheximide and paromomycin, lead to more distinct phenotypes for wildtype and *trm2*Δ strains. Relative to wildtype, *trm2*Δ cells were more sensitive to cycloheximide, while they were less sensitive to hygromycin B and paromomycin **(Figure 2A and Extended Data Figure 1)**. Growth assays in media concur with our spot plating observations (**Supplemental Figure S1**), and measurement of the minimum inhibitory concentration of hygromycin B for *trm2*Δ and Δ*trmA S. cerevisiae* and *E. coli*, respectively, revealed that m^5^U54-deplete cells are less sensitive to the drug **(Figure 2B and Extended Data Figure 2)**. We elected to continue follow-up studies with cycloheximide and hygromycin B because we have found that multiple cell lines lacking a variety of different tRNA modifying enzymes are strongly desensitized to paromomycin, but the cycloheximide and hygromycin B were more *trm2*Δ and Δ*trmA* specific. Consistent with this observation, RNA-seq reveals that the transcriptome profile of *trm2*Δ cells is less disrupted under hygromycin B stress relative to wild-type cells (**Figure 3A-D**), and the expression of a luciferase reporter in the presence of hygromycin B is increased *trm2*Δ cells **(Figure 2C)**.

**Figure 2.**
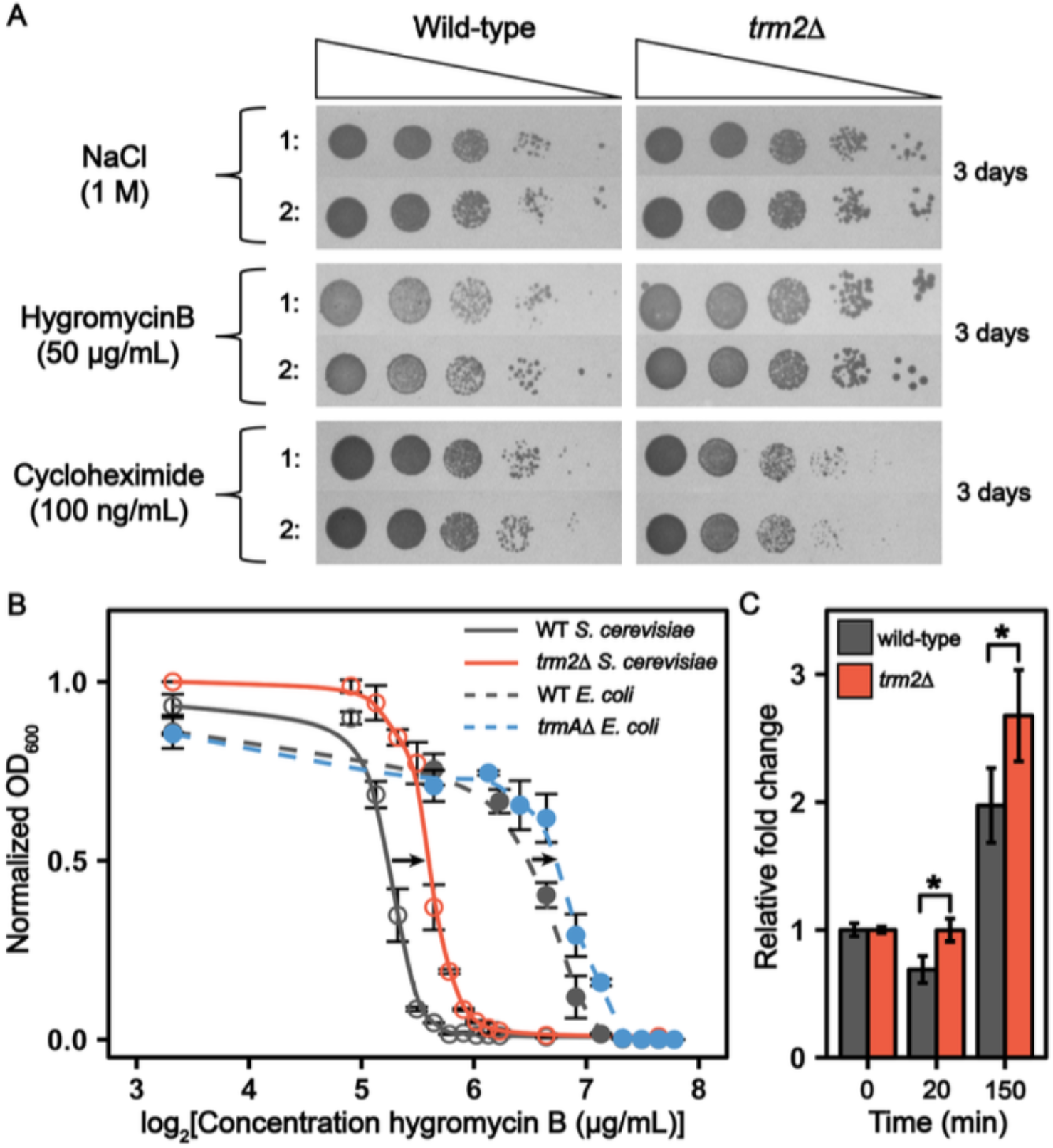
*trm2*Δ *S. cerevisiae* modulate cell growth under cellular stress. (A) Spot platting assays comparing cellular growth between WT and *trm2*Δ BY4741 *S. cerevisiae* with NaCl, hygromycin B and cycloheximide. **(B)** Hygromycin B minimum inhibitory concentration curves for WT BY4741 *S. cerevisiae*, *trm2*Δ BY4741 *S. cerevisiae*, WT *E. coli* K-12 BW25113, and Δ*trmA E. coli* K-12 BW25113. The OD_600_ for each hygromycin B concentration was recorded after 24 hr growth and normalized to the growth without antibiotic. **(C)** Luciferase reporter assay displaying the luciferase protein produced after the addition of 50 μg/mL hygromycin B. Both cell lines were grown to mid-log phase (OD600 = 0.5) prior to the addition of hygromycin B. Luminescence was measured at multiple times following drug addition (0 min, 20 min, 60 min, 120 min), and increased in the *trm2*Δ cells compared to the wildtype after hygromycin B was added.

**Figure 3.**
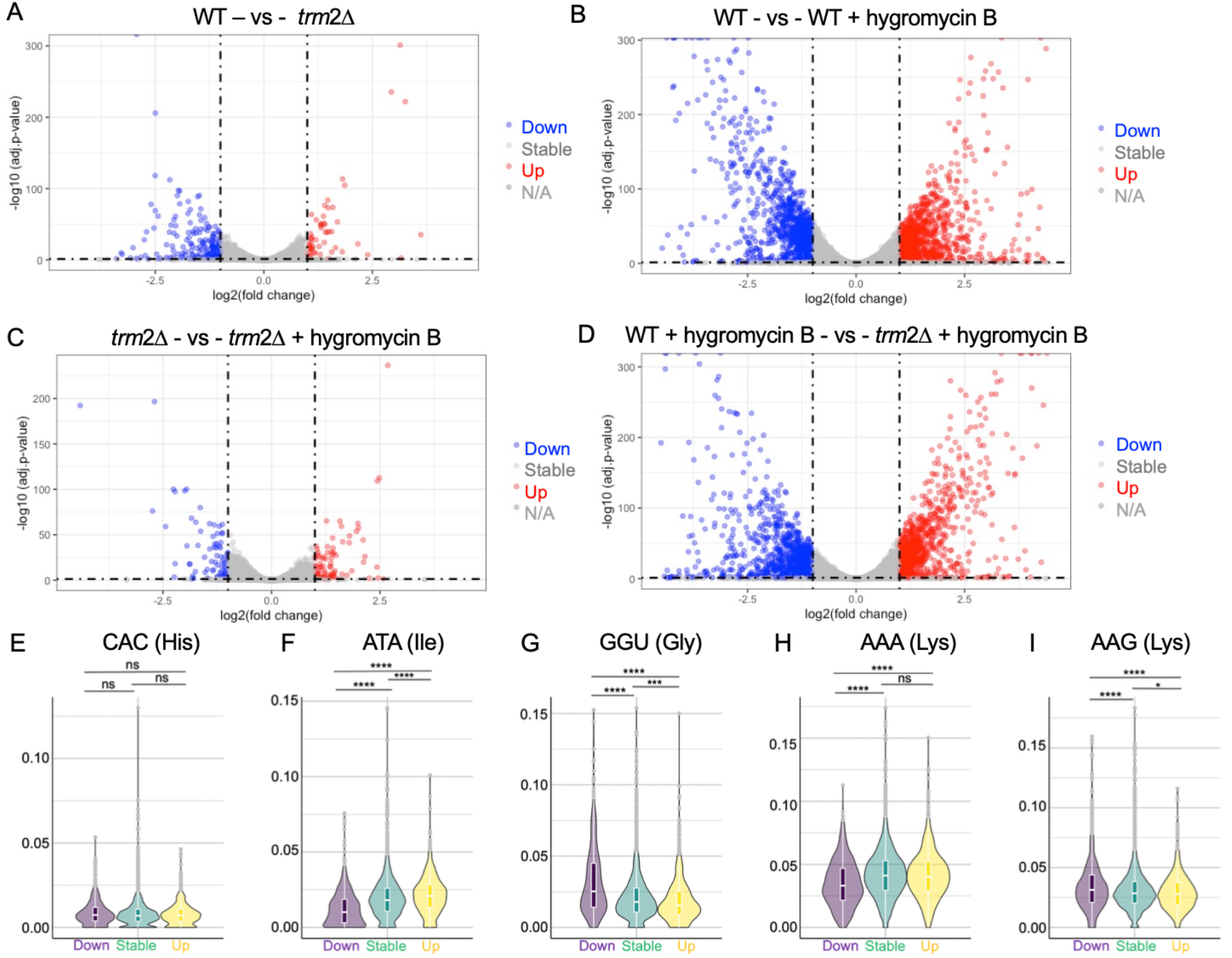
mRNA gene expression is less impacted in trm2Δ than wildtype *S. cerevisiae* following hygromycin B treatment. Log_2_ folded change of the mean of normalized counts for each transcript detected by RNA-seq for: (**A**) wild-type vs *trm2*Δ *S. cerevisiae* grown in unstressed (YPD, no hygromycin B) conditions; (**B**) wild-type *S. cerevisiae* untreated with hygromycin B vs wild-type *S. cerevisiae* treated with hygromycin B; (**C**) *trm2*Δ untreated with hygromycin B vs *trm2*Δ treated with hygromycin B; (**D**) wild-type vs *trm2*Δ *S. cerevisiae* treated with hygromycin B.; (**E-I**) Log_2_ folded change of the mean of normalized counts for each transcript detected by RNA-seq for: (**A**) wild-type vs *trm2*Δ *S. cerevisiae* grown in unstressed (YPD, no hygromycin B) conditions; (**B**) wild-type *S. cerevisiae* untreated with hygromycin B vs wild-type *S. cerevisiae* treated with hygromycin B; (**C**) *trm2*Δ untreated with hygromycin B vs *trm2*Δ treated with hygromycin B; (**D**) wild-type vs *trm2*Δ *S. cerevisiae* treated with hygromycin B.; (**E-I**) Usage distribution of the indicated codon in genes with RNA levels that go down (purple), remain stable (green) or increase (yellow) in WT cells treated with hygromycin B. Codon usage is normalized to gene length.

### mRNA m^5^U modification does not drive TRM2-dependent changes in cellular fitness

In yeast m^5^U is incorporated into both tRNAs and mRNAs^19,20,22^. Therefore, the TRM2-dependent differences in cellular fitness that we observed in the presence of hygromycin B could arise from changes to the modification landscape of either RNA species. We implemented a multiplexed UHPLC-MS/MS assay developed in our lab to measure the levels of 50 different modifications within mRNAs isolated from untreated, hygromycin B- and cycloheximide-treated wild-type cells^19^. Total mRNA was purified using a previously described three-stage purification pipeline that treats total RNA to small RNA depletion, rRNA depletion and two rounds of poly(A) RNA isolation^19^. Subsequently, mRNA purity was confirmed using Bioanalyzer, RNA-seq, RT-qPCR, and LC-MS/MS **(Extended Data Figure 3 and Supplemental Figures S2 and S3)**. We find the levels of a limited subset mRNA modifications increase (e.g., pseudouridine, 2’O-methylguanosine) or decrease (e.g., m^5^U) modestly (≤ 1.5 -fold) in response to sub-MIC concentrations of hygromycin B and cycloheximide (**Figure 4A**). The levels of m^5^U in mRNAs isolated from antibiotic treated cells is extremely low (0.003 ± 0.001 % m^5^U/U in hygromycin B and 0.0025 ± 0.0006 % m^5^U/U in cycloheximide) and the modification is unlikely to be regularly encountered by the ribosome as there is only 1 m^5^U substitution per every ∼35,000 Us (**Supplemental Table 1**). Taken together with our previous *in vitro* translation studies demonstrating that the inclusion of m^5^U into mRNA codons does not slow the peptide elongation significantly^19^, our data suggest that the TRM2-dependent effects on cell growth that we observe do not originate from the action of the enzyme on mRNAs. Instead, they indicate that differential impact of hygromycin B on wild-type and *trm2*Δ on cellular fitness and protein production more likely results from the changes in m^5^U54-tRNA modification status.

**Figure 4.**
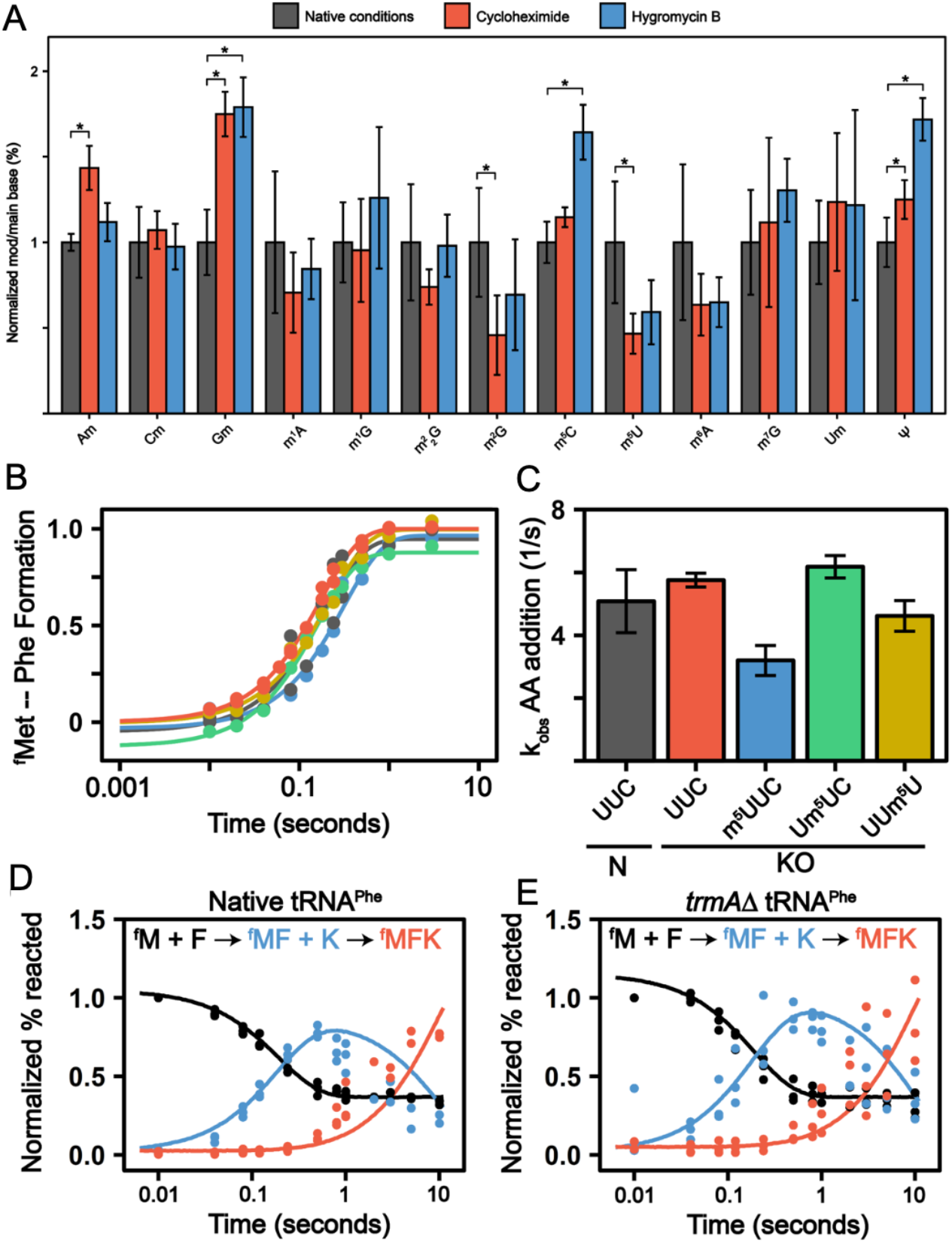
(A) Cycloheximide and hygromycin B translational inhibition modestly alters mRNA modification landscape. Normalized modification abundance with purified mRNA (modification/main base %) in WT BY4741 *S. cerevisiae* grown without antibiotic (grey), 100 ng/mL cycloheximide (red), or 50 μg/mL hygromycin B (blue). Error bars correspond to the standard deviation of two biological replicates. * corresponds to a significant alteration where p < 0.05. **(B)** k_obs_ curves and **(C)** values for Met-Phe dipeptide formation using the tRNA^Phe,-m5U^. **(D & E)** Time courses displaying the formation of ^f^Met-Phe-Lys tripeptide with 1 μM Phe+Lys TC complex using either **(D)** WT *E. coli* purified tRNA^Phe^ or **(E)** Δ*trmA E.coli* purified tRNA^Phe^ with WT *E.coli* purified tRNA^Lys^.

### m^5^U54 in tRNA^Phe^ does not influence the rate constant for amino acid addition

The changed sensitivity of the *trm2*Δ knockout strains for growth in sub-MIC concentrations of the translation elongation inhibitors raises the possibility that m^5^U54 modulates the elongation step in the protein synthesis pathway. We implemented a well-established fully reconstituted *E. coli in vitro* translation system was used to directly test this supposition^25^. This system has long been used to conduct high-resolution kinetic studies investigating how the ribosome decodes mRNAs. The core mechanism of translation elongation is well conserved between bacteria and eukaryotes^26,27^, and prior studies demonstrate that mRNA modifications that slow elongation and/or change mRNA decoding elongation in the reconstituted *E. coli* also do so in eukaryotes^28,29^. We first evaluated if m^5^U54 influences the rate of amino acid addition by comparing the rate constants for a single amino acid addition to a growing polypeptide using *E. coli* tRNA purified from either wild type (tRNA^Phe^) or Δ*trmA* cell lines (tRNA^Phe,-m5U^) **(Figures 4B-C)**.

In our assays, *E. coli* 70S ribosome initiation complexes were formed on mRNAs encoding a ^f^Met-Phe-Lys tri-peptide, with ^35^S-labeled ^f^Met-tRNA^fMet^ bound in the ribosome P site. Initiation complexes were reacted with an excess of ternary complex containing a single aminoacylated tRNA (Phe-tRNA^Phe^ or Phe-tRNA^Phe,-m5U^), elongation factor Tu (EF-Tu) and GTP. The formation of ^35^S-^f^Met-Phe dipeptide product is followed over time to determine the observed rate constant (*k*_obs_) for Phe addition. tRNA^Phe,-m5U^ was aminoacylated as efficiently as tRNA^Phe^ by an excess of recombinant *E. coli* phenylalanine tRNA synthetase. Consistent with previous reports, the rate constants for Phe incorporation that we measured are comparable for tRNA^Phe^ and tRNA^Phe,-m5U^ (*k*_obs_ ∼ 5s^-1^; **Figure 4B-C**).

Additionally, we investigated the possibility of cooperative affects between m^5^U containing mRNA codons and m^5^U54 in tRNA^19–22^. To accomplish this, we interrogated how amino acid addition on m^5^U-containing codons (1^st^, 2^nd^, and 3^rd^ position modified UUU or UUC codons) is impacted when decoded by tRNA^Phe,-m5U^. These findings were compared with assays we previously published investigating amino acid addition on m^5^U-modified codons using a fully modified tRNA^Phe19^. As on the unmodified codons, the rate constants for amino acid addition were essentially unchanged when tRNA^Phe^ and tRNA^Phe,-m5U^ were used in translation assays **(Figure 4C)**^19^. Our data collectively demonstrate that loss of m^5^U54 in tRNA^Phe,-m5U^ does not alter the overall rate constant for Phe addition on unmodified or m^5^U modified codons.

### Loss of m^5^U54 in P site tRNA^Phe^ enables ribosome translocation in the presence of hygromycin B

Hygromycin B blocks translation by interacting with the RNA in the ribosome A site to prevent translocation^30,31^. In the single amino acid addition assays described above the ribosome does not need to translocate to form the ^f^Met-Phe di-peptide. As such, these assays do not report on the step in protein synthesis for which we have a cellular phenotype (**Figure 2**). Therefore, we next examined the impact of m^5^U54 by following the formation of ^f^Met-Phe-Lys tri-peptide, which does require ribosome translocation to occur. In these experiments, we treated the 70S initiation complexes with ternary complexes that included either EF-Tu:Phe-tRNA^Phe^:GTP or Phe-tRNA^Phe,-m5U^, EF-Tu:Lys-tRNA^Lys^:GTP and the elongation factor, EFG, required for efficient ribosome translocation. In the absence of hygromycin B, ^f^Met-Phe-Lys synthesis robustly occurred with both Phe-tRNA^Phe^ and Phe-tRNA^Phe,-m5U^ **(Figures 4D-E)**. When hygromycin B was included in the reaction mix the formation of the ^f^Met-Phe dipeptide was still rapid with tRNA^Phe^ and tRNA^Phe,-m5U^ **(Figure 5A)**. However, as expected, no ^f^Met-Phe-Lys formation is observed in the presence of hygromycin B in assays performed with tRNA^Phe^. In contrast, ^f^Met-Phe-Lys is generated when tRNA^Phe,-m5U^ is used instead, in line with our observation that *trm2*Δ *S. cerevisiae* cells produce more reporter protein than wildtype cells in the presence of hygromycin B **(Figure 2C)**. These data suggest that inclusion of m^5^U54 in tRNA^Phe^ impacts the step in the elongation cycle between peptidyl transfer and subsequent aa-tRNA binding, translocation.

**Figure 5.**
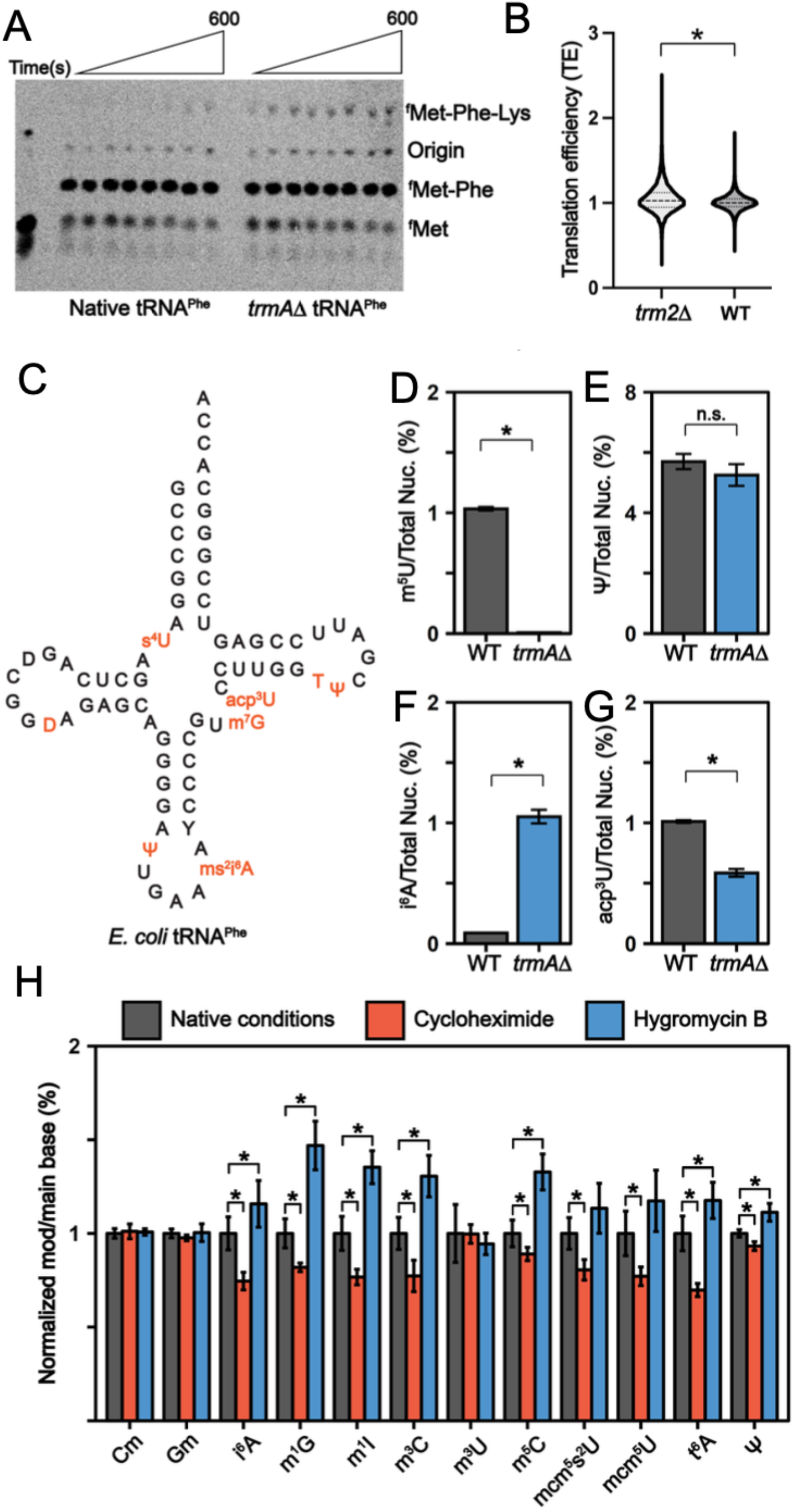
TRM2 and TRMA enhance translation and tRNA modification. (A) Δ*trmA E. coli* purified tRNA^Phe^ improve tripeptide synthesis in the presence of hygromycin B inhibition. eTLC displaying the formation of ^f^Met-Phe-Lys tripeptide with 1 μM Phe+Lys TC complex in the presence of 50 μg/mL hygromycin B using either (Left) WT *E. coli* purified tRNA^Phe^ or (Right) Δ*trmA E. coli* purified tRNA^Phe^ with WT *E. coli* purified tRNA^Lys^. **(B)** Translation efficiency (TE) in wildtype and *trm2*Δ *S. cerevisiae*. TEs are replotted from Chou, et al.^39^ (**C-H**) RNA modification landscape fluctuates. **(C)** WT *E. coli* tRNA^Phe^ sequence containing known modifications. By LC-MS/MS, the modification abundance (modification/total nucleoside %) was determined for WT and Δ*trmA E. coli* purified tRNA^Phe^ for **(D)** m^5^U, **(E)** Ψ, **(F)** i^6^A, and **(G)** acp^3^U. *corresponds to a significant alteration where p < 0.05. We estimate the i^6^A stoichiometry be 0.8 modifications per tRNA^Phe,-m5U^. **(H)** Normalized modification abundance (modification/main base %) in WT BY4741 *S. cerevisiae* grown without antibiotic (grey), 100 ng/mL cycloheximide (red), or 50 μg/mL hygromycin B (blue) for modifications found in stem loop of *S. cerevisiae* tRNA.

#### TRMA modulates the modification landscape of tRNA^Phe^

It is becoming apparent that “modification circuits” exist in tRNAs, and a hierarchy has been observed for the installation of modifications in the T loop region of yeast tRNAs, with m^5^U54 positively influencing the introduction of m^1^A58^32,33^. This led us consider that loss of m^5^U54 in Δ*trmA E. coli* might change the stoichiometry of other tRNA modifications. To test this possibility, we used LC-MS/MS to quantitatively measure the levels of the m^5^U, 4-thiouridine (s^4^U), dihydrouridine (D), pseudouridine (Ψ), 7-methylguanosine (m^7^G), and 3-(3-amino-3-carboxypropyl)uridine (acp^3^U) modifications in tRNA^Phe^ purified from wild-type and 11*trmA E. coli* (tRNA^Phe,-m5U^). Additionally, we examined the levels the N6-isopentenyladenosine (i^6^A) component of the 2-methylthio-N6-isopentenyladenosine (ms^2^i^6^) modification, as we were unable to acquire a sufficiently soluble nucleoside standard required to directly measure ms^2^i^6^A levels in our multiplexed LC-MS/MS approach. We find that m^5^U54 is completely lost from tRNA^Phe,-m5U^, as expected (**Figure 5D**). Furthermore, while most other tRNA^Phe^ modifications were unaltered (**Figure 5E and Extended Data Figure 4)**, the levels of two modifications did change. There was a moderate decrease in the inclusion of the variable loop modification acp^3^U (1.7-fold, **Figure 5G)** and a significant increase in i^6^A (12-fold, **Figure 5F)**, consistent with observations made using orthogonal approaches (*Schultz, et al*)^34^. We posit that the increased level of i^6^A likely results from the depletion of the ms^2^i^6^A37 hyper-modification. Thus, it appears that, like its yeast homolog TRM2, the action of TRMA influences the maturation of sites on tRNAs beyond m^5^U54.

#### S. cerevisiae tRNA anticodon stem loop modification landscape is impacted by antibiotic treatment

While tRNA modifications have long been thought to be stoichiometric and statically incorporated, evidence is emerging that many modifications are incorporated at sub-stoichiometric levels and that changes in tRNA modification status can modulate the sensitivity of cells anti-fungal, anti-cancer and anti-bacterial drugs^35–38^. Our dual observations that the loss of TRMA changes the modification profile of *E. coli* tRNA^Phe^, and that lack of 5-methyluridine transferases in both bacteria (Δ*trmA*) and yeast (*trm2*Δ) alter the sensitivity of cells to antibiotics led us to wonder if the translational inhibitors that we investigated here impact the tRNA modification landscape. To test this, we again turned to our highly sensitive multiplexed UHPLC-MS/MS assay to measure the level of 50 RNA modifications in total RNA isolated from *S. cerevisiae* treated with cycloheximide and hygromycin B. These studies revealed that levels of tRNA-specific modifications, but not rRNA-specific modifications (e.g., m^3^U), fluctuated in response to the antibiotics (**Figure 5H and Extended Data Figure 5**). The largest effects were observed for modifications incorporated in tRNA anticodon stem loops at position 37 (e.g. m^1^I, t^6^A, i^6^A and m^1^G) (**Figure 5H and Extended Data Figure 5**). These findings correlate with our observation that the distribution of codons decoded by tRNAs with modifications at position 37 show changes in RNA levels upon hygromycin B (**Figure 3E-F and Supplemental Figure S4**). In general, cycloheximide and hygromycin B treatment had opposite effects on modification incorporation - with cycloheximide reducing modification levels, and hygromycin B promoting their inclusion. The most significant differences between conditions were seen for i^6^A, t^6^A, m^1^I and m^1^G - which are roughly twice as prevalent in RNAs isolated from hygromycin treated cells than those treated with cycloheximide. All four of these modifications (and their related derivative modifications) are well-established modulators of ribosome elongation speed and fidelity^39^.

## DISCUSSION

There are few RNA modifications as highly conserved across biology as m^5^U54 is in tRNAs. The universal addition of this modification is a long-standing mystery, as the enzymes that incorporate m^5^U54 are not essential, and previous studies suggest that loss of m^5^U54 does not affect the step in translation that most tRNAs participate in, elongation^14–17^. To address this conundrum, we employed genetic methods to study translation in cells lacking this modification, and *in vitro* biochemistry using tRNAs without m^5^U54 in their TΨC loop. We find that m^5^U54 restricts translocation *in vitro*, consistent with published data indicating that the loss of m^5^U54 subtly enhances translation efficiency *in vitro* and in cells (**Figures 2 and 5B**)^39,40^. Furthermore, we reveal that the efficacy of the translocation inhibitors hygromycin B and cycloheximide is altered in strains lacking m^5^U54 (**Figure 2**). Our data provide evidence that m^5^U54 influences gene expression, as hygromycin B induces substantial changes in the mRNAs present in wild-type cells that are alleviated by loss of the m^5^U54 modifying enzyme (*trm2*Δ) (compare **Figure 3B and 3C)**. Finally, our studies reveal that m^5^U54 installation is critical for maintaining the global tRNA landscape, significantly promoting the inclusion of modifications in the anticodon stem loop (e.g. ms^2^i^6^A, t^6^A, m^1^I; **Figure 5C-H and Extended Data Figure 4**). Previous observations indicate that m^5^U54 limits the dynamics of the TΨC loop, suggesting that tRNAs lacking m^5^U54 are more flexible than their fully modified counterparts^23,41^. Together with our findings, this suggests a model wherein m^5^U54 rigidifies the TΨC loop to maintain the critical link between mRNA and peptide sequence, at the cost of decreased cellular fitness under some environmental stresses.

As discussed above, our data indicate that one biological consequence of m^5^U54 is the modulation of translocation, both directly and indirectly (through subtly altering the tRNA modification landscape). In the protein synthesis pathway, translocation takes place following the transfer of the growing polypeptide chain from the P site peptidyl-tRNA to an aminoacyl-tRNA bound in the ribosome A site. Elongation factor G (EFG) is responsible for catalyzing the movement of the ribosome to reposition the A site peptidyl-tRNA in the P site. Hygromycin B blocks this process by flipping a key 16S rRNA nucleotide (A1493) between the A and P sites^30,42,43^. Increased flexibility in the TΨC loop of m^5^U54-deplete tRNAs may provide a satisfying molecular rationale for our observation that *trm2*Δ and tRNA^Phe,-m5U^ partially overcome hygromycin B inhibition in cells and *in vitro* (**Figures 2, 3 and 4A)**. Such a model suggests that in unstressed circumstances m^5^U54 modulates translocation, and can sometimes slow the process, as we observed with tRNA^Phe,-m5U^ (**Figure 6**). This is consistent with tRNAs is consistent long-perplexing findings from Nierhaus and co-workers that lysine peptides are generated more quickly *in vitro* from poly(A) transcripts using m^5^U54-deplete tRNAs, but that neither tRNA binding or peptidyl-transfer are affected^40^. Similarly, recent observations indicate that total protein production is increased from reporters containing select amino acid repeats (e.g. Leu(TTA), Tyr(TAT), Ile(ATA)) in a Δ*trmA E. coli* strain (*Schultz, submitted*)^34^. Lastly, comparison of available ribosome profiling for wild-type and *trm2*Δ yeast supports the supposition that m^5^U54 can subtly alter translation; there is a small, but statistically significant, global increase in the translational efficiency of ribosomes in *trm2A*11 cells (**Figure 5B**)^39^.

**Figure 6:**
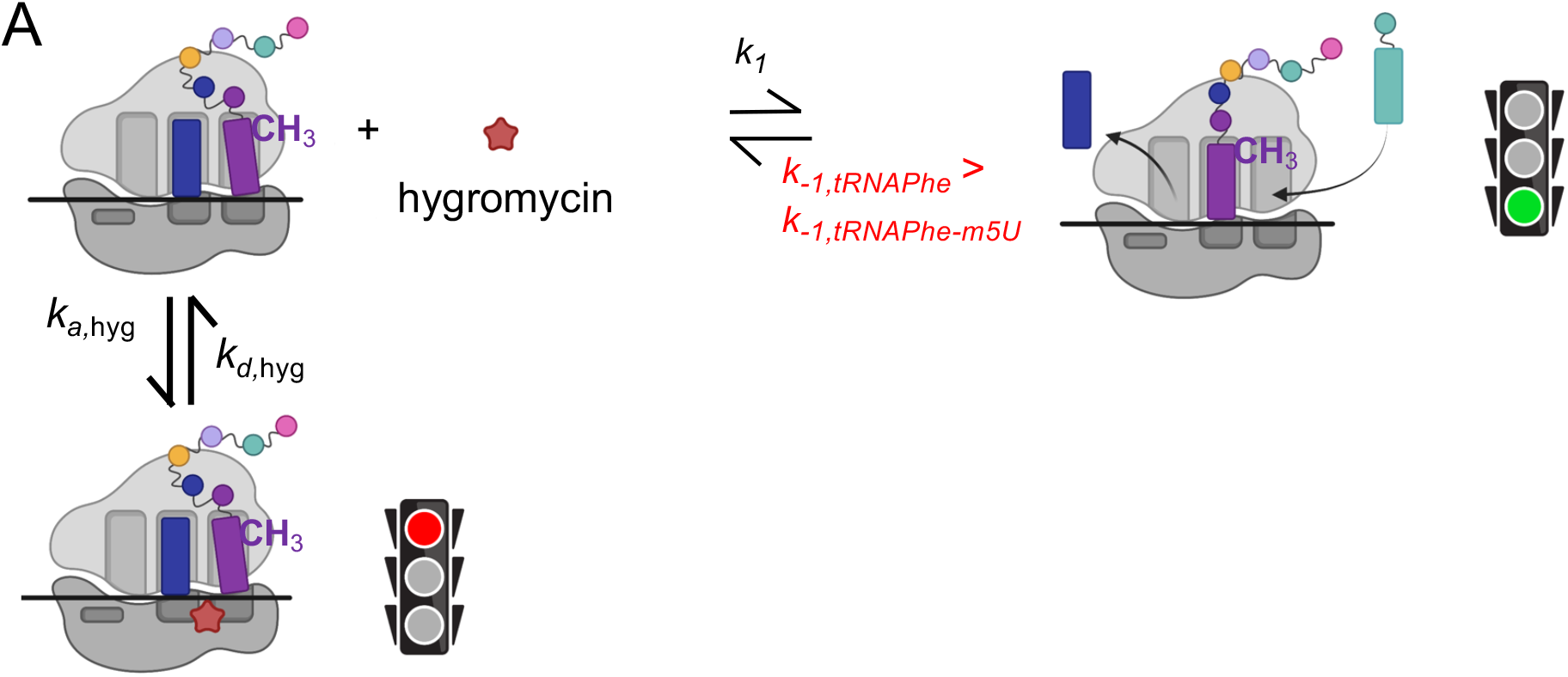
Model for mitigation of hygromycin B sensitivity by tRNA^Phe,-m5U^. (A) Hygromycin B acts to block translocation, using a non-competitive mechanism to flip ribosome nucleotides and prevent movement of the A site peptidyl-tRNA into the P site. In this mechanism, when translation takes place with natively modified tRNAs that include m^5^U54 (panel A), the observed rate constant for hygromycin B binding is much faster than that of translocation leading to translational stalling (*k*_obs, hyg_ >> *k*_obs, translocation_). However, when m^5^U54 is missing from tRNAs the rate constant for translocation (*k*_1_) is increased sufficiently to raise the *k*_obs, translocation_ such that it now effectively competes with the k_obs, hyg_, allowing for some translocation to occur and product to be generated (k_obs, hyg_ ∼ k_obs, translocation_). Given the levels of peptide observed in our assays, *k*_obs, translocation_ shifts to be within 10-fold of k_obs, hyg_ when reactions are performed with tRNA^Phe-m5U^.

This model lead to us to wonder how the negative modulation of translational efficiency and translocation might be beneficial given that wild-type cells outcompete m^5^U54 depleted strains (*trm2*Δ and Δ*trmA*) ^16^. Recent computational studies indicate that differences in the local dynamics of the TΨC loop in cognate and near cognate tRNAs during accommodation act as gatekeepers for ensuring accurate mRNA decoding^44^. We speculate that perturbed tRNA dynamics may increase flux through a non-canonical pathway tRNA accommodation pathway that is independent of codon:anticodon interactions^44^. Needless to say, increased decoupling peptide sequence from mRNA sequence is likely to be detrimental to cells under optimal growth conditions. Additionally, by controlling translocation rates, m^5^U4 also has the potential to increase the accuracy of translation by limiting potentially deleterious processes such as frameshifting reliant on translocation^45,46^.

The implication of m^5^U54 in translocation is reasonable give that the elbow region of ribosome-bound tRNAs (formed by tertiary interactions between the TΨC- and D-loops) mediate their coordination with the ribosome^47^. Therefore, residues in the elbow are likely to be critically important for efficient translocation. Consistent with this, the mutation of tRNA positions adjacent to m^5^U54 (Ψ55 and C56) decrease the translocation rate^48^. Notably, recent Cryo-EM snapshots taken throughout translocation reveal that elbow moves sufficiently to alter the bend of the peptidyl-tRNA by up to 17.37° during translocation, further supporting the notion that tRNA elbow dynamics are critical^47^. Our studies add to the idea that the TΨC plays a crucial role in translocation, suggesting that m^5^U54 helps to maintain the tRNA elbow region shape and dynamics necessary for mediating tRNA interactions with the ribosome during translocation.

Not all of the subtle changes in translation observed in cells lacking m^5^U54 likely arise from m^5^U54 directly tweaking translocation. Our LC-MS/MS analyses of tRNA modification landscapes indicate that m^5^U54 installation is correlated to alterations in the modification status of other tRNA positions, including acp^3^U and ms^2^i^6^A in *E. coli* tRNA^Phe^ (**Figure 5F-G**). These results agree with the findings of MSR-tRNA sequencing studies by Kothe, et al. demonstrating that acp^3^U, ms^2^i^6^A and s^4^U levels are altered in several (but not all) tRNAs, including tRNA^Phe^, in *ΔtrmA* cells (*Schultz, submitted*)^34^. Presumably, as not all tRNAs possess the same set of modifications, tRNA-specific changes may arise that have differential impacts on translocation.

The seemingly small changes in translocation or tRNA modification stoichiometry that we observe upon the loss of m^5^U54 can have an outsized impact on gene expression under stress conditions. Our work reveals that under stress TRM2 helps to shape the mRNA landscape (**Figure 3**), supporting the idea that m^5^U54 alters translational efficiency given that translation rates are tied to mRNA stability^39^. This is in line with recent studies suggesting that translocation, despite not being the overall rate limiting step for protein synthesis, never the less can impact protein production^47,49^. Notably, the hygromycin B dependent alterations in wildtype yeast gene mRNA levels that we observe appear to be codon dependent (**Figure 3E-F and Supplementary Figure S4**), consistent with codon-dependent effects of TRMA on translation recently reported in *E. coli* (*Schultz, submitted*)^34^. Although the effects that we observed on MIC were admittedly modest, the decreased sensitivity of *trm2*Δ to hygromycin B augments the growing body of literature suggesting that tRNA modifying enzymes play significant roles in altering gene expression to modulate the development of drug resistance^37^.

Collectively, our findings address the decades long question of why m^5^U54 is conserved across tRNAs in all organisms, identifying multiple roles for the modification in the protein synthesis pathway. Our LC-MS/MS analyses reveal a that loss of the enzyme responsible m^5^U54 installation alters the tRNA modification landscape in both *S. cerevisiae* and *E. coli*. Additionally, kinetic and cell-based assays provide evidence that m^5^U54 serves to modulate the translocation step in protein synthesis, to ultimately impact gene expression (especially when cells are under translational stress). Our data support structural work revealing that the tRNA TΨC stem loop-forms critical interactions with the ribosome during translocation, and demonstrate how even apparently small perturbations in the modification profiles and inherent dynamic landscape of tRNAs is important enough to drive the conservation of TΨC modifications throughout biology. As we seek to assign biological functions for other RNA modifications in the mechanism protein synthesis, the findings of this study suggest that even relatively subtle changes in the dynamic processes the drive translation can are likely significant in the nature, when cells perform under non-optimized (non-laboratory) conditions.

## METHODS

### Spot plating growth assay and growth curve characterization under stress

Wild-type and *trm2*Δ cells were inoculated into 3 mL YPD and grown overnight. These cultures were diluted to OD_600_=1, and 7 μl of 10-fold serial dilutions were spotted on fresh YPD agar plates including 0.75-1.0 M NaCl, 250 mM MgSO_4_, 200 μM puromycin, 100 ng/mL cycloheximide, 25-50 μg/mL hygromycin B, 50 μM MG132 and 1.5-3 mg/mL paromomycin. Growth of the cells were also tested in the presence of different carbon sources including 2% glucose, 2% sucrose, 2% galactose and 3% glycerol in YEP agar media (1% *S. cerevisiae* extract and 2% peptone). The plates were incubated for 2-5 days at 30 °C unless indicated.

For growth curves, wild-type and *trm2Δ* cells were inoculated into 5 mL YPD and grown overnight. Cultures were then diluted to a starting OD_600_ = 0.05 – 0.1 in 100 mL YPD media containing either 1 M NaCl, 0.1 µg/mL cycloheximide, 50 µg/mL hygromycin B, or 3 mg/mL paromomycin. Cultures were grown in duplicate at 30°C with shaking unless indicated, and growth was monitored by OD_600_.

### Minimum inhibitory concentration of hygromycin B in *S. cerevisiae* and *E. coli*

Wildtype BY4741 and *trm2*Δ BY4741 (Dharmacon) *S. cerevisiae* were inoculated in 5 mL of YPD and grown overnight at 30℃ shaking at 285 RPM. Overnight cultures were diluted to 0.02 OD_600_ in YPD containing various concentrations of hygromycin B. The cells were cultured for 24 hr at 30℃ shaking at 285 RPM. The growth was measured by OD_600_.

The *E. coli* Keio knockout parental strain BW25113 (Dharmacon) and Δ*trmA E. coli* JW3937 (Dharmacon) were inoculated in 5 mL of LB and grown overnight at 37℃ shaking at 250 RPM. Overnight cultures were diluted to 0.002 OD_600_ in LB containing various concentrations of hygromycin B. The cells were cultured for 24 hr at 37℃ shaking at 250 RPM. The growth was measured by OD_600_.

### Luciferase reporter assay

Plasmid was transformed into wild-type and *Δtrm2 S. cerevisiae* by the standard lithium-acetate/PEG protocol. The cells were streaked onto CSM-URA agar plates to isolate single colonies^50^. CSM-URA media (30 mL) was inoculated with a single colony and allowed to grow overnight at 30℃ and 250 RPM. The cells were diluted to an OD_600_ of 0.05 with 500 mL of CSM-URA medium and were grown to an OD_600_ of 0.5 at 30℃ and 250 RPM. At this point, the cells were stressed with hygromycin B or cycloheximide and were allowed to continue to grow. At time points of 0 min, 20 min, 60 min, and 150 min after the translational stress, 10 mL of culture was pelleted at 8,000 x g for 10 min. The cell pellet was washed with 1 mL of water prior to storage at −80℃ until the assay was performed.

The cell pellets were washed with 500 μL of breaking buffer (50 mM sodium phosphate pH 7.4, 2 mM EDTA, 10% glycerol, 2 mM PMSF) prior to resuspension in 200 μL of breaking buffer. An equal volume of acid washed glass beads were added, and the tubes were vortexed for 30 seconds followed by 30 seconds on ice. The vortexing and ice incubation was repeated three times for a total of four times. The lysates were centrifuged for 21,000 x g for 15 min at 4℃. The supernatant (30 μL per replicated) was characterized by the Promega dual luciferase assay kit.

### *S. cerevisiae* cell growth and mRNA purification

Wild-type BY4741 and *Δtrm2 S. cerevisiae* (Dharmacon) were grown in YPD medium as previously described^51^. *Δtrm2 Saccharomyces cerevisiae* were grown in the presence of 200 μg/mL Geneticin. Briefly, 10 mL of YPD medium was inoculated with a single colony selected from a plate and allowed to grow overnight at 30℃ and 250 RPM. The cells were diluted to an OD_600_ of 0.1 with 200 mL of YPD medium and were grown to an OD_600_ between 0.6 and 0.8 at 30℃ and 250 RPM. Translational stress *S. cerevisiae* were grown with 50 μg/mL hygromycin B or 100 ng/mL cycloheximide. Hygromycin B *S. cerevisiae* were grown to an OD_600_ of 0.4 to ensure cells were in mid-log phase growth. This cell culture was pelleted at 15,000 x g at 4℃ and used for the RNA extraction.

*S. cerevisiae* cells were lysed as previously described with minor alterations^19^. The 200 mL cell pellet was resuspended in 8 mL of lysis buffer (60 mM sodium acetate pH 5.5, 8.4 mM EDTA) and 800 μL of 10% SDS. One volume (8.8 mL) of phenol was added and vigorously vortexed. The mixture was incubated at 65℃ for five minutes and was again vigorously vortexed. The incubation at 65℃ and vortexing was repeated once. Then, the mixture was rapidly chilled in an ethanol/dry ice bath and centrifuged for 15 minutes at 15,000 x g. The total RNA was extracted from the upper aqueous phase using a standard acid phenol-chloroform extraction. The extracted total RNA was treated with 140 U RNase-free DNase I (Roche, 10 U/μL) at 37℃ for 30 min. The DNase I was removed through an acid phenol-chloroform extraction. The resulting total RNA was used for our UHPLC-MS/MS, bioanalyzer, and RNA-seq analyses.

mRNA was purified through a three-step purification pipeline^19^. First, small RNA (tRNA and small rRNA) was diminished using the MEGAclear Transcription Clean-Up Kit (Invitrogen) to purify RNA >200nt. Then, two consecutive Dynabeads oligo-dT magnetic bead selections (Invitrogen, USA) were used to purify poly(A) RNAs from 140 μg of small RNA depleted RNA. The resulting RNA was ethanol precipitated and resuspended in 14 μL. Subsequently, we used the commercial riboPOOL rRNA depletion kit (siTOOLs Biotech, Germany) to remove residual 5S, 5.8S, 18S, and 28S rRNA. The Bioanalyzer RNA 6000 Pico Kit (Agilent, USA) was used to evaluate the purity of the mRNA prior to UHPLC-MS/MS analysis.

### qRT-PCR

The RevertAid First Strand cDNA Synthesis Kit (Thermo Scientific, USA) was used to reverse transcribe DNase I-treated total RNA and three-stage purified mRNA (200 ng) using the random hexamer primer. The resulting cDNA was diluted 5,000-fold and 1 μL of the resulting mixture was analyzed using the Luminaris Color HiGreen qPCR Master Mix (Thermo Scientific, USA) with gene-specific primers^19^.

### RNA -seq

The WT *S. cerevisiae* total RNA and mRNA were analyzed by RNA-seq as previously described by paired-end sequencing using 2.5% of an Illumina NovaSeq (S4) 300 cycle sequencing platform flow cell (0.625% of flow cell for each sample)^19^. All sequence data are paired-end 150 bp reads. The reads were trimmed using Cutadapt v2.3^52^. The reads were evaluated with FastQC^53^ (v0.11.8) to determine quality of the data. Reads were mapped to the reference genome Saccharomyces_cerevisiae (ENSEMBL), using STAR v2.7.8a and assigned count estimates to genes with RSEM v1.3.3^54,55^. Alignment options followed ENCODE standards for RNA-seq (Dobin). QC metrics from several different steps in the pipeline were aggregated by multiQC v1.7^56^. Differential expression analysis was performed using DESeq2 ^57^. Codon usage was calculated using. coRdon: Codon Usage Analysis and Prediction of Gene Expressivity. R package version 1.18.0, https://github.com/BioinfoHR/coRdon)^58^.

### RNA enzymatic digestion and UHPLC-MS/MS ribonucleoside analysis

Total RNA and mRNA (125 ng) were digested for each condition. The RNA was hydrolyzed to composite mononucleosides using a two-step enzymatic reaction and quantified using LC-MS/MS as previously described with no alterations^19^.

### *E. coli* ribosomes, and translation factors tRNA and mRNA for in vitro assay

Ribosomes were purified from *E. coli* MRE600 as previously described^28^. All constructs for translation factors were provided by the Green lab unless specifically stated otherwise. Expression and purification of translation factors were carried out as previously described^28^. Unmodified transcripts were prepared using run-off transcription of a DNA template. Modified mRNA sequences containing m^5^U were purchased HPLC purified from Dharmacon. The mRNA sequence used was 5’-GGUGUCUUGCGAGGAUAAGUGCAUU<u>AUGUUCUAA</u>GCCCUUCUGUAGCCA-3’, with the coding sequence underlined. The m^5^U modified position was always the first position in the UUC phenylalanine codon.

Native *E. coli* tRNA^Phe^ was purified as previously described with minor alterations^25^. Bulk *E. coli* transfer RNA was purified in *E. coli* form an HB101 strain containing pUC57-tRNAphe provided by Yury Polikanov. Two liters of enriched Terrific Broth (TB) media (TB, 4 mL glycerol/L, 50 mM NH_4_Cl, 2 mM MgSO_4_, 0.1 mM FeCl_3_, 0.05% glucose and 0.2% lactose (if autoinduction media was used)) were inoculated with 1:400 dilution of a saturated overnight culture and incubated at 37°C overnight shaking at 250 RPM with 400 mg/mL of ampicillin. Cells were harvested the next morning by 30 min centrifugation at 5000 RPM and then stored at −80℃ until lysis was performed. To lyse the cells, the pellet was resuspended in 200 mL of resuspension buffer (20mM Tris-Cl, 20 mM Mg(OAc)_2_ pH 7) and 100 mL of phenol:chloroform:isoamyl alcohol pH 4.5 (125:24:1) was added. The mixtures were incubated at 4℃ shaking at 250 RPM for 1 hr. After incubation, the lysate was centrifuged at 3,220×g for 60 min at 4℃, and the aqueous supernatant was transferred to a fresh tube. The phenol:chloroform:isoamyl alcohol was washed with 100 mL of 18 MΩ water, the mixture was centrifuged at 3,220×g for 60 min at 4℃, and the aqueous supernatant was transferred to the same tube. The pooled aqueous supernatant was washed with 100 mL chloroform, centrifuged at 3,220×g for 60 min at 4℃, and the supernatant was transferred to a fresh tube. DNA was precipitated with 150 mM NaOAc pH 5.2 and 20% isopropanol. The DNA was removed through centrifugation at 13,700×g for 60 minutes at 4℃. The supernatant was transferred to a fresh tube, the isopropanol content was raised to 60%, and the bottle was stored at −20℃ to precipitate the RNA. The precipitation was centrifuged at 13,700×g for 60 minutes at 4℃ and the supernatant was removed. The pellet was resuspended in 10 mL 200 mM tris-acetate pH 8.0 and incubated at 37℃ for 2 hr. After incubation, the RNA was precipitated through the addition of 1/10^th^ volume of 3 M NaOAC pH 5.2 and 2.5 volumes of 100% ethanol. The precipitate could either be stored at −20℃ or centrifuged immediately at 16,000×g for 60 min at 4℃. The pellet was washed with 70% ethanol, resuspended in 5-10 mL of water, and desalted using a 3K MWCO Amicon centrifugal concentrator.

Total tRNA was isolated by anion exchange chromatography on a FPLC using a Cytiva Resource Q column. The mobile phase A was 50 mM NH_4_OAc, 300 mM NaCl, 10 mM MgCl_2_ and mobile phase B was 50 mM NH_4_OAc, 800 mM NaCl, 10 mM MgCl_2_. The resuspend RNA was injected (5 mL) and eluted with a linear gradient from 0% to 50% B over 18 column volumes. Fractions were pulled and ethanol precipitated overnight at −20℃.

The RNA was centrifuged at 16,000×g for 30 min at 4℃, washed with 70% ethanol, and resuspended in approximately 500 μL of water. tRNA^Phe^ was purified from total tRNA using a Waters XBridge BEH C18 OBD Semiprep column (10 x 250 mm, 300 Å, 5 μm). Mobile phase A was 20 mM NH_4_OAc, 10 mM MgCl_2_, 400 mM NaCl at pH 5 and mobile phase B was 20 mM NH_4_OAc, 10 mM MgCl_2_, 400 mM NaCl at pH 5 with 60% methanol. The total tRNA was injected (400 μL) and separated with a linear gradient of buffer B from 0-35% was done over 35 minutes. The MPB% was increased to 100% over 5 min and held at 100% MPB for 10 min. The column was then equilibrated for 10 column volumes before next injection at 0% MPA. Each fraction was analyzed by A_260_ absorbance and amino acid acceptor activity to identify fractions that contain tRNA^Phe^.

### Formation of *E. coli* ribosome initiation complexes

Initiation complexes (IC’s) were formed in 1X 219-Tris buffer (50 mM Tris pH 7.5, 70 mM NH_4_Cl, 30 mM KCl, 7 mM MgCl_2_, 5 mM ß-ME) with 1 mM GTP as previously described^25^. 70S ribosomes were incubated with 1 μM mRNA (with or without modification), initiation factors (1,2,3) all at 2 μM final and 2 μM of radiolabeled ^f^Met-tRNA for 30 min at 37℃. After incubation MgCl_2_ was added to a final concentration of 12 mM. The ribosome mixture was then layered onto 1 mL cold buffer D (20 mM Tris-Cl, 1.1 M sucrose, 500 mM NH_4_Cl, 10 mM MgCl_2_, 0.5 mM disodium EDTA, pH 7.5) and centrifuged at 69,000 rpm for 2 hr at 4℃. After pelleting, the supernatant was discarded into radioactive waste, and the pellet was resuspended in 1X 219-tris buffer and stored at −80℃.

### *In vitro* amino acid addition assays: dipeptide formation

Initiation complexes were diluted to 140 nM with 1X 219-tris buffer. Ternary complexes (TCs) were formed by first pre-loading EF-Tu with GTP (1X 219-tris buffer, 10 mM GTP, 60 μM EFTu, 1 μM EFTs) at 37℃ for 10 min. The EF-Tu mixture was incubated with the tRNA mixture (1X 219-tris buffer, Phe-tRNA^Phe^ (1-10 μM), 1 mM GTP) for another 15 min at 37℃. After TC formation was complete, equal volumes of IC complexes (70 nM) and ternary complex (1 μM) were mixed either by hand or using a KinTek quench-flow apparatus. Discrete time-points (0-600 seconds) were taken as to obtain observed rate constants on m^5^U-containing mRNAs. Each time point was quenched with 500 mM KOH (final concentration). Time points were then separated by electrophoretic TLC and visualized using phosphorimaging as previously described ^25,28^. Images were quantified with ImageQuant. The data were fit using Equation 1:

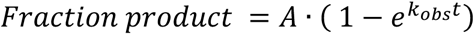

### *In vitro* translation amino acid addition assays for tripeptide formation

Initiation complexes were diluted to 140 nM with 1X 219-Tris buffer. Ternary complexes (TCs) were formed by first pre-loading EF-Tu with GTP (1X 219-tris buffer, 10 mM GTP, 60 μM EFTu, 1 μM EFTs) at 37℃ for 10 min. The EF-Tu mixture was incubated with the tRNA mixture (2 μM aminoacyl-tRNA Phe/Lys(s), 24 μM EF-G, 60 μM EF-Tu) with ICs (140 nM) in 219-Tris buffer (50 mM Tris pH 7.5, 70 mM NH_4_Cl, 30 mM KCl, 7 mM MgCl_2_, 5 mM βME) for 15 min at 37℃. These experiments are done with both native phenylalanine tRNA or our Δm5U phenylalanine tRNA. After TC formation was complete, equal volumes of IC complexes (70 nM) and ternary complex (1 μM) were mixed using a KinTek quench-flow apparatus. Discrete time-points (0-600 seconds) were taken to obtain observed rate constants on non-modified mRNAs, containing a UUC phenylalanine codon. Each time point was quenched with 500 mM KOH (final concentration). Time points were then separated by electrophoretic TLC and visualized using phosphorimaging as previously described^25,28^. Images were quantified with ImageQuant. The data were fit using Equation 1 as previously described.

### *In vitro* translation amino acid addition assays for tripeptide formation in the presence of Hygromycin B

Initiation complexes were diluted to 140 nM with 1X 219-tris buffer. Ternary complexes (TCs) were formed by first pre-loading EF-Tu with GTP (1X 219-tris buffer, 10 mM GTP, 120 μM EFTu, PEP, 12mM, PK .40μM, 40 μM EFTs) at 37℃ for 10 min. The EF-Tu mixture was incubated with the tRNA mixture (20–60 μM aminoacyl-tRNA Phe/Lys(s), 24 μM EF-G, 60 μM EF-Tu) with ICs (140 nM) in 219-tris buffer (50 mM Tris pH 7.5, 70 mM NH_4_Cl, 30 mM KCl, 7 mM MgCl_2_, 5 mM βME) for another 15 min at 37℃. These experiments are done with both native phenylalanine tRNA or our Δm5U phenylalanine tRNA. After TC formation was complete, 50μg/ml of Hygromycin B was added to the IC complex. Then by hand equal volumes of IC complexes (70 nM) and ternary complex (1 μM) were mixed and discrete time-points (0-600 seconds) were taken as to obtain observed rate constants on non-modified mRNAs, containing a UUC phenylalanine codon. Each time point was quenched with 500 mM KOH (final concentration). Time points were then separated by electrophoretic TLC and visualized using phosphorimaging as previously described ^25,28^. Images were quantified with ImageQuant. The data were fit using Equation 1 as previously described.

### Fitting observed rate constants and global analysis simulations of amino acid addition

Fits to obtain observed rate constant *k_1_* was done using a single differential equation in KaleidaGraph for products in di peptide formation. When multiple peptide products were formed, the disappearance of ^f^Met product was fit using a single exponential equation in KaleidaGraph to get an observed rate constant *k_1_*. This value was then used in KinTex Explorer to measure subsequent rate constant *k_2_* using simulations. Simulations were modeled against the equation:

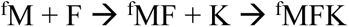

## ACKNOWLEDGEMENTS

We thank the National Institutes of Health (NIGMS R35 GM128836 to KSK, 4R00GM135533-03 to RON, and T32 GM132046 to MFK) and National Science Foundation (CAREER Award 2045562 to KSK, CHE-1904146 to RTK, and GRFP to JDJ) for their support. Additionally, we are grateful for insight provided by discussions with Dr. Shura Mankin.

## Extended Data

**Extended Data Figure 1:**
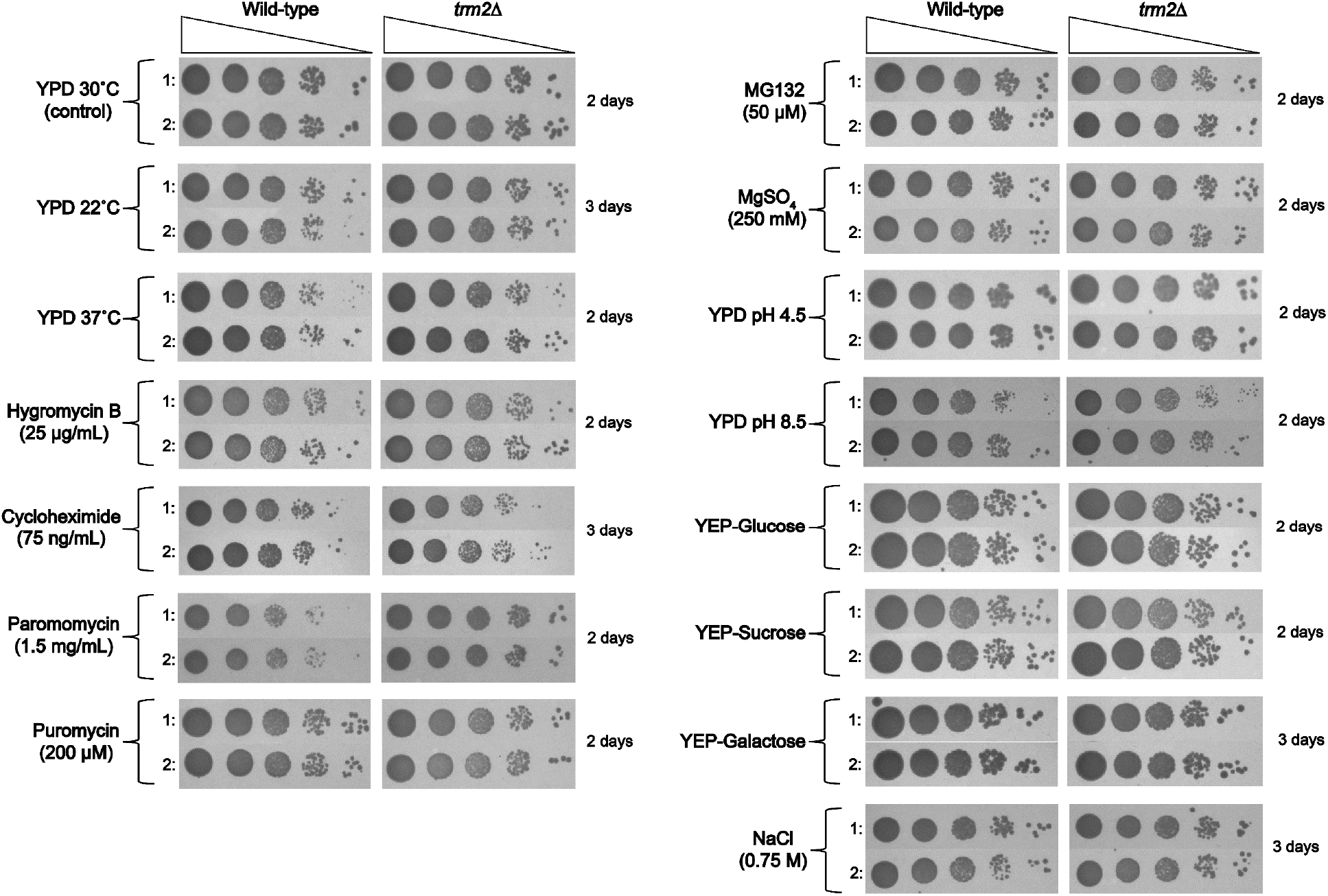
Spot plating analysis for WT and *trm2*Δ BY4741 *S. cerevisiae* under cellular stress conditions. The cell growth of WT and *trm2*Δ BY4741 *S. cerevisiae* were compared under multiple cellular stress conditions. Unless otherwise stated, the agar plates were incubated at 30℃ for the specified amount of time.

**Extended Data Figure 2:**
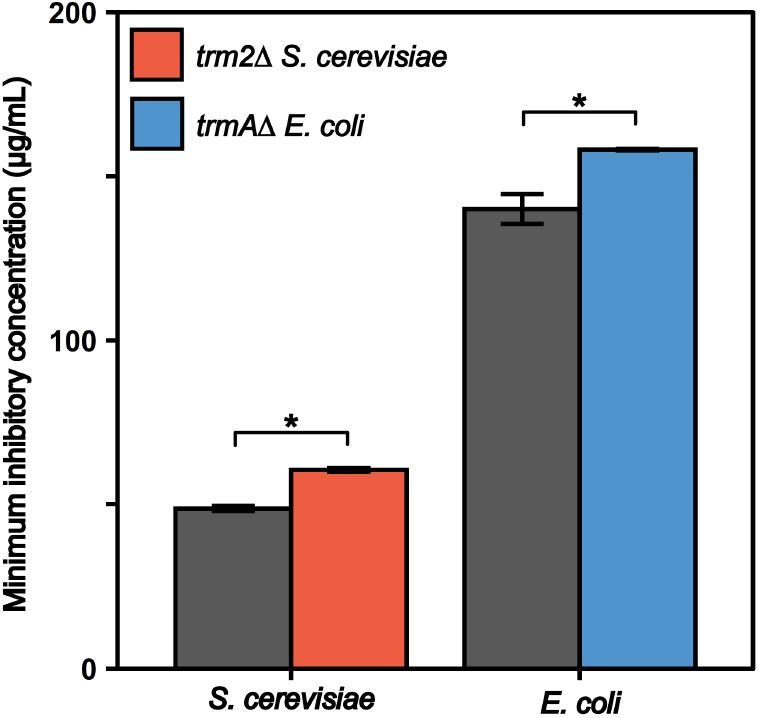
*E. coli* and *S. cerevisiae* lacking tRNA (uracil-5-)-methyltransferase are less susceptible to hygromycin B inhibition. The hygromycin B minimum inhibitory concentration for WT and knockout *E. coli* and *S. cerevisiae* are provided. The error bars correspond to the standard deviation between two biological replicates. * corresponds to a significant alteration where p < 0.05.

**Extended Data Figure 3:**
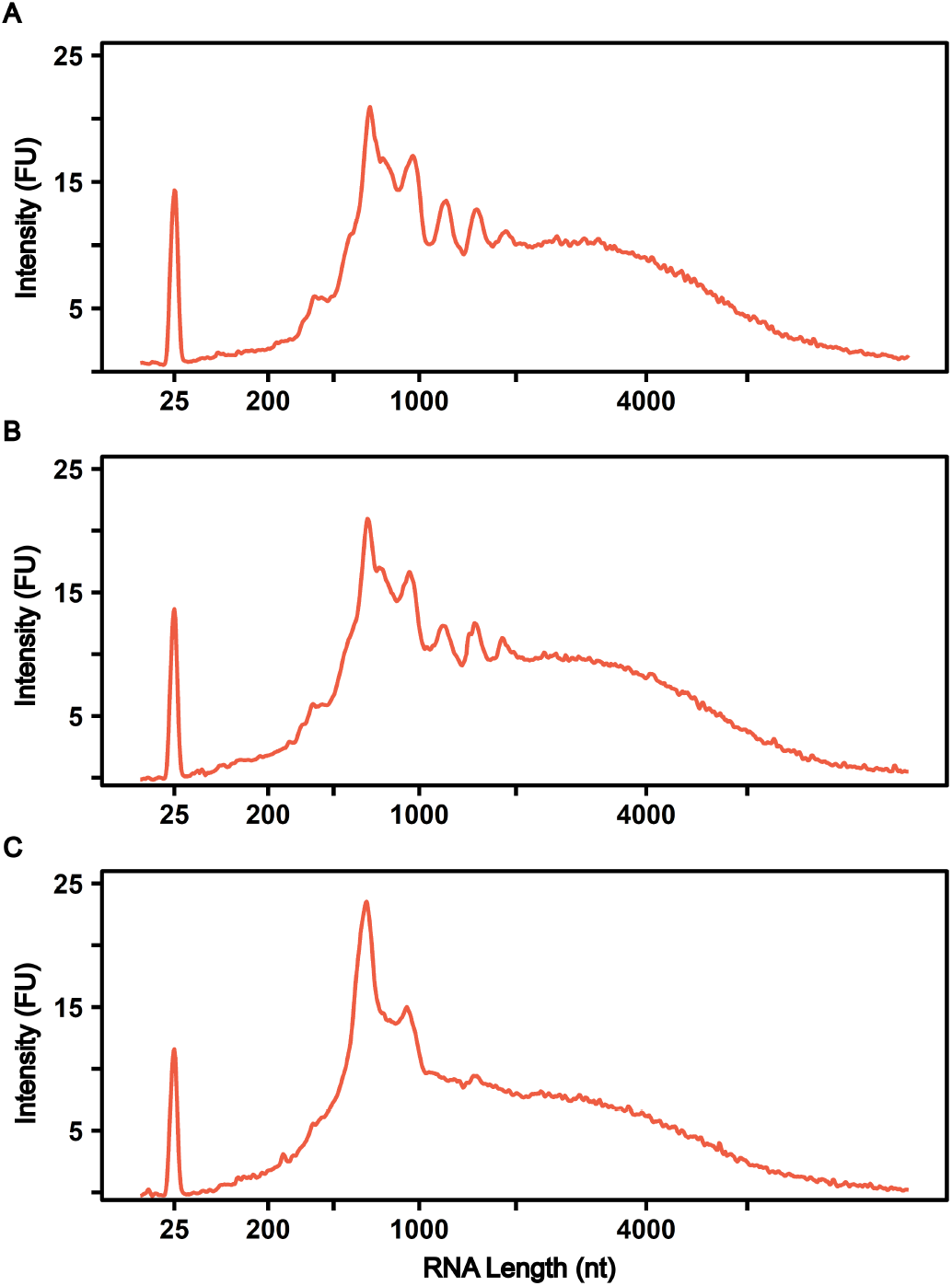
Bioanalyzer traces of mRNAs purified from yeast cells for LC-MS/MS analysis. Representative Bioanalyzer electropherograms of purified mRNA from BY4741 *S. cerevisiae* grown **(A)** without translational inhibitor, **(B)** 100 ng/mL cycloheximide, or **(C)** 50 μg/mL hygromycin B.

**Extended Data Figure 4:**
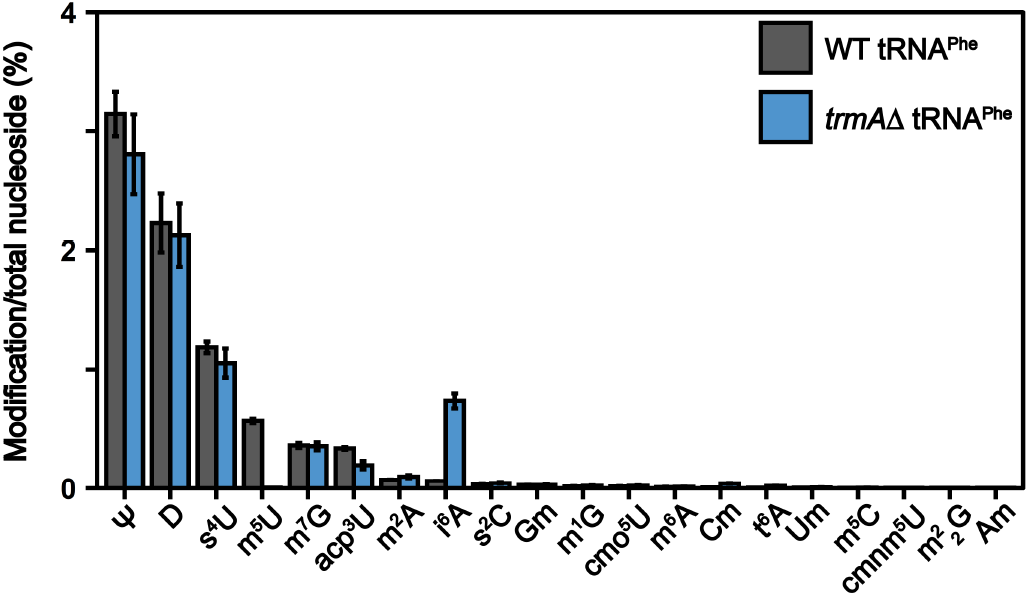
tRNA^Phe^ from *E. coli* lacking tRNA (uracil-5-)-methyltransferase has altered modification landscape. Modification abundance (modification/total nucleoside %) determined by LC-MS/MS in tRNA^Phe^ purified from WT and Δ*trmA E. coli*.

**Extended Data Figure 5:**
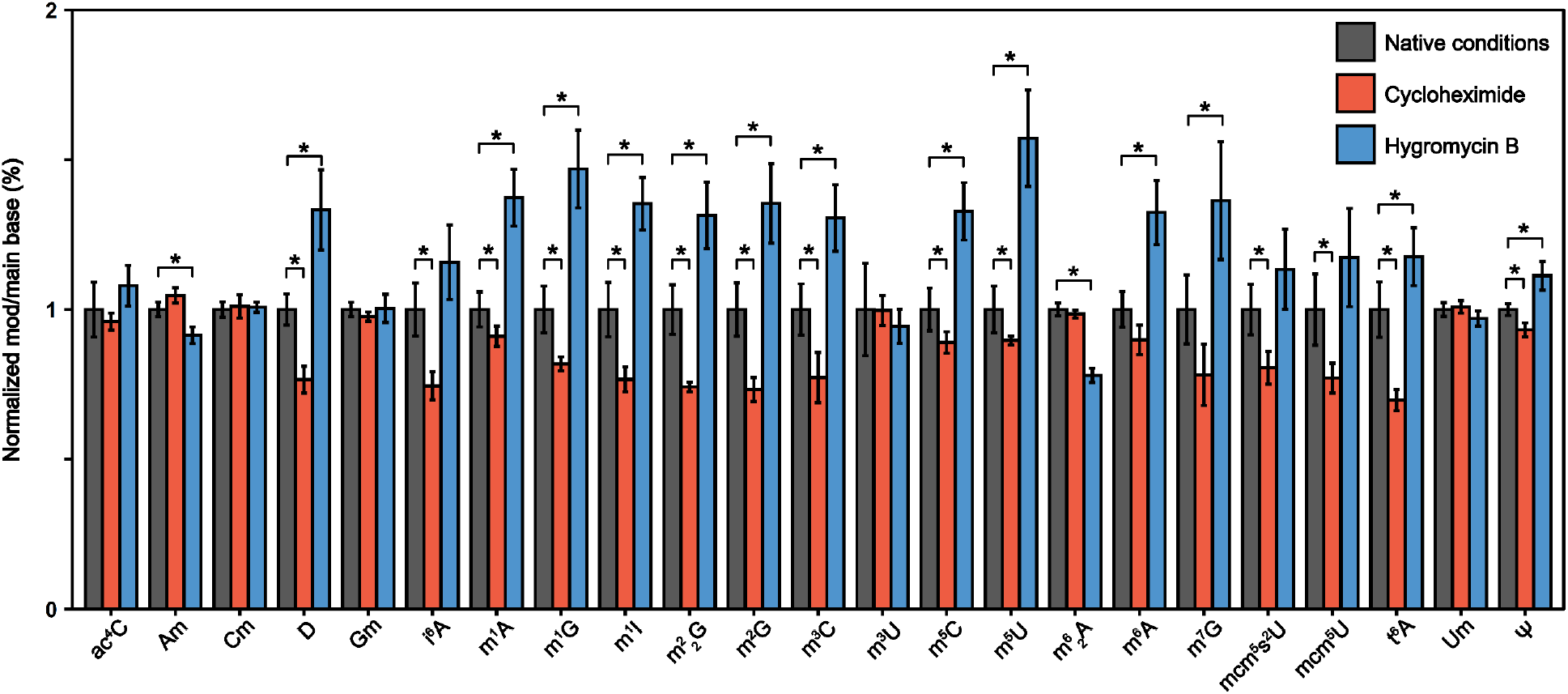
Cycloheximide and hygromycin B translational inhibition alters ncRNA modification landscape. Normalized modification abundance (modification/main base%) in WT BY4741 *S. cerevisiae* grown without antibiotic (grey), 100 ng/mL cycloheximide (red), or 50 μg/mL hygromycin B (blue). Error bars correspond to the standard deviation of two biological replicates. * corresponds to a significant alteration where p < 0.05.

## Notes

### Competing Interest Statement

The authors have declared no competing interest.

