## Supplemental Figures for "Conserved 5-methyluridine tRNA modification modulates ribosome translocation"

OD600

| Time (min) | WT - NaCl #1 | dTrm2 - NaCl #1 | WT - NaCl #2 | dTrm2 - NaCl #2 | WT - CHX #1 | dTrm2 - CHX #1 | WT - CHX #2 | dTrm2 - CHX #2 | WT - Hygromycin #1 | dTrm2 - Hygromycin #1 | WT - Hygromycin #2 | dTrm2 - Hygromycin #2 |
| --- | --- | --- | --- | --- | --- | --- | --- | --- | --- | --- | --- | --- |
| 815 | 0.33 | 0.42 | 0.33 | 0.38 | 0.53 | 0.43 | 0.51 | 0.43 | 0.24 | 0.44 | 0.26 | 0.38 |
| 875 | 0.39 | 0.49 | 0.39 | 0.45 | 0.62 | 0.51 | 0.59 | 0.51 | 0.27 | 0.48 | 0.27 | 0.41 |
| 935 | 0.46 | 0.58 | 0.45 | 0.54 | 0.73 | 0.59 | 0.69 | 0.6 | 0.28 | 0.51 | 0.27 | 0.42 |
| 995 | 0.53 | 0.67 | 0.52 | 0.62 | 0.82 | 0.67 | 0.8 | 0.68 | 0.29 | 0.51 | 0.28 | 0.43 |
| 1055 | 0.6 | 0.73 | 0.59 | 0.68 | 0.93 | 0.78 | 0.89 | 0.76 | 0.26 | 0.5 | 0.27 | 0.42 |
| 1115 | 0.69 | 0.89 | 0.68 | 0.81 | 1.08 | 0.91 | 1.06 | 0.93 | 0.32 | 0.55 | 0.31 | 0.47 |
| 1175 | 0.86 | 1.38 | 0.86 | 1.26 | 1.58 | 1.4 | 1.64 | 1.42 | 0.37 | 0.62 | 0.36 | 0.51 |
| 1235 | 1.28 | 1.6 | 1.36 | 1.48 | 2.32 | 2.04 | 2.28 | 2.04 | 0.33 | 0.61 | 0.33 | 0.48 |
| 1295 | 1.44 | 2.08 | 1.48 | 1.92 | 2.8 | 2.32 | 2.6 | 2.24 | 0.34 | 0.6 | 0.34 | 0.52 |
| 1355 | 1.84 | 2.36 | 1.84 | 2.24 | 3.36 | 2.76 | 3.2 | 2.76 | 0.37 | 0.67 | 0.33 | 0.53 |
| 1415 | 1.96 | 2.68 | 1.92 | 2.32 | 3.8 | 3 | 3.5 | 3.08 | 0.37 | 0.68 | 0.33 | 0.53 |
| 1460 | 2.25 | 2.95 | 2.2 | 2.85 | 4.1 | 3.5 | 3.9 | 3.4 | 0.35 | 0.62 | 0.34 | 0.52 |
| 1490 | 2.3 | 3.1 | 2.2 | 2.85 | 4.3 | 3.6 | 4.25 | 3.65 | 0.34 | 0.61 | 0.32 | 0.5 |

**Supplemental Figure S1: *trm2*Δ *S. cerevisiae* display cellular growth phenotype under cellular stress.** Cellular growth times courses for wildtype and *trm2*Δ *S. cerevisiae* under osmotic (NaCl) stress, cycloheximide (CHX) stress, and hygromycin stress for two biological replicates. At each time point an OD_600_ measurement was collected.

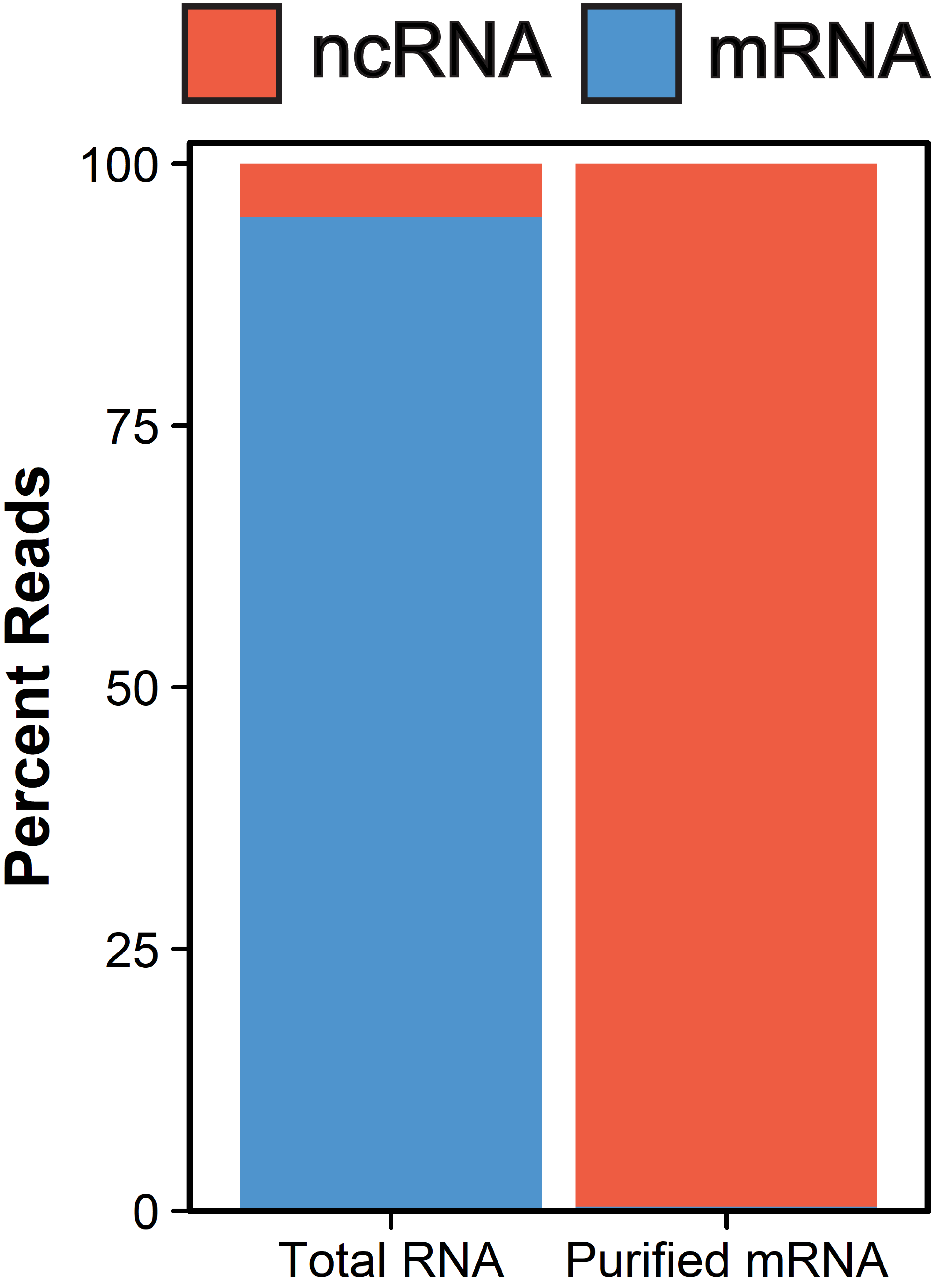

**Supplemental Figure S2: RNA-seq confirms high purity of *S. cerevisiae* mRNA.** The percent of ncRNA (red) and mRNA (blue) reads within *S. cerevisiae* total RNA and purified mRNA are displayed.

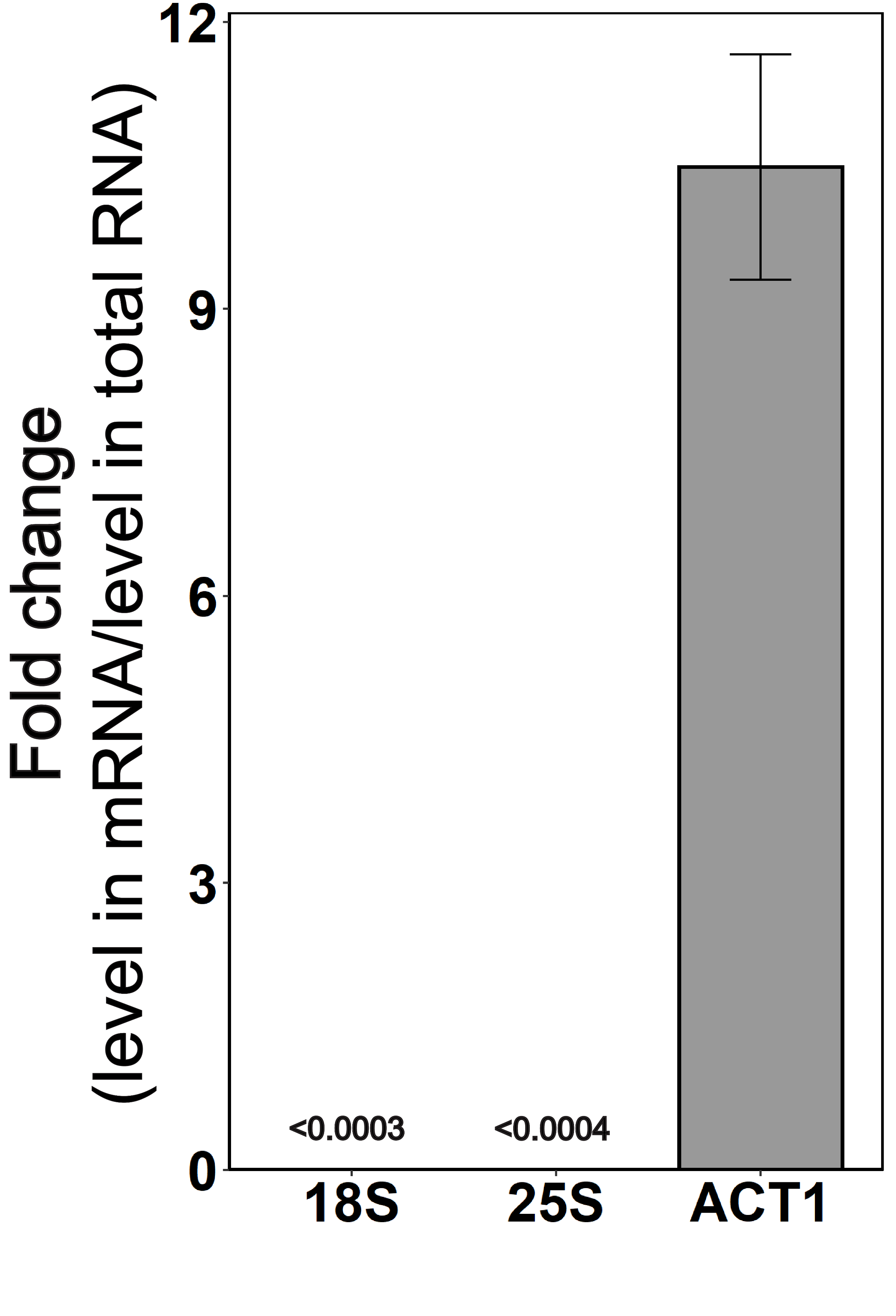

**Supplemental Figure S3: RT-qPCR displays enrichment of mRNA in purified mRNA.** Fold change in purified mRNA to total RNA for the *S. cerevisiae* 18S rRNA, 25S rRNA, and ACT1 mRNA.

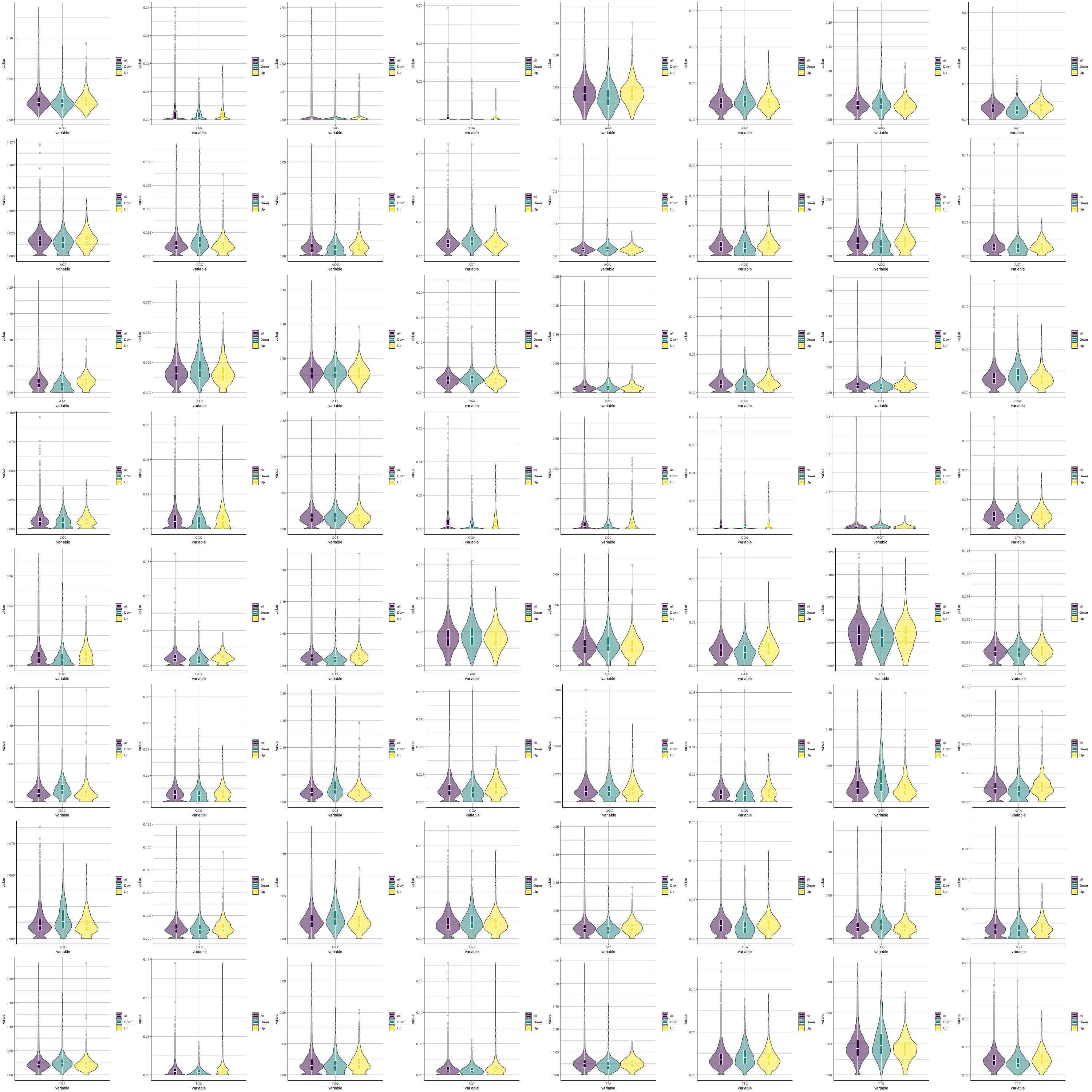

**Supplemental Figure S4:** Usage distribution of the indicated codon in genes with overall RNA levels (purple), and those that go down (green), or increase (yellow) in WT cells treated with hygromycin B. Codon usage is normalized to gene length.
